# A Microphysiological Model of Progressive Human Hepatic Insulin Resistance

**DOI:** 10.1101/2025.01.08.631261

**Authors:** Dominick J. Hellen, Jessica Ungerleider, Erin Tevonian, Pierre Sphabmixay, Priyatanu Roy, Caroline Lewis, Jacob Jeppesen, Damien Demozay, Linda G. Griffith

## Abstract

**Background & Aims:** Hepatic insulin resistance is a fundamental phenomenon observed in both Type 2 diabetes (T2D) and metabolic (dysfunction) associated fatty liver disease (MAFLD). The relative contributions of nutrients, hyperinsulinemia, hormones, inflammation, and other cues are difficult to parse *in vivo* as they are convoluted by interplay between the local and systemic events. Here, we used a well-established human liver microphysiological system (MPS) to establish a physiologically-relevant insulin-responsive metabolic baseline and probe how primary human hepatocytes respond to controlled perturbations in insulin, glucose, and free fatty acids (FFAs).

**Methods:** Replicate liver MPS were maintained in media with either 200 pM (normal) or 800 pM (T2D) insulin for up to 3 weeks. Conditions of standard glucose (5.5 mM), hyperglycemia (11 mM glucose), normal (20µM) and elevated FFA (100 µM), alone and in combination were used at each insulin concentration, either continuously or reversing back to standard media after 2 weeks of simulated T2D conditions. Hepatic glucose production, activation of signaling pathways, insulin clearance, transcriptome analysis, and intracellular lipid and bile acid accumulation were assessed.

**Results:** Hyperinsulinemia alone induces insulin resistance after one week of exposure, while hyperglycemia and increased FFAs significantly exacerbate this phenotype. Hyperinsulinemia, along with elevated glucose and FFAs, transcriptionally predisposes hepatocytes to insulin resistance through altered metabolic and immune signaling pathways. The phenotypes observed in hyperinsulinemia and nutrient overload are partially reversible upon return to normophysiologic conditions.

**Conclusions:** Our enhanced *in vitro* model, replicating multiple aspects of the insulin-resistant condition, offers improved insights into disease mechanisms with relevance to human physiology.

**Lay Summary:** The many nutritional, hormonal, and systemic inflammatory factors that contribute to the loss of sensitivity to insulin in Type 2 Diabetes and other metabolic disorders are difficult to parse in human patients, and animal models fail to capture all the human dimensions of the relevant biology. We, therefore, developed a microphysiological systems model, involving a microfluidic platform that cultures a simulated human liver for weeks at a time, under controlled nutrient and hormone concentrations, to analyze how the effects of nutrition (glucose and free fatty acids) together with increased insulin cause pathological features seen in human liver *in vivo*. We found that continually high levels of insulin predispose the liver to some of these features, along with reversal when the nutritional and insulin levels were restored to healthy values.

## Introduction

Type 2 diabetes (T2D) and metabolic (dysfunction) associated fatty liver disease (MAFLD) are chronic diseases affecting millions of individuals worldwide. ^1^ Over 70% of T2D patients also suffer from MALFD with high incidence of metabolic associated steatohepatitis (MASH). ^1^ In addition to excess fat accumulation in the liver, a pathophysiological hallmark of MAFLD, insulin resistance in liver, muscle, and adipose tissue are retained and augmented by chronic inflammation as MAFLD progresses into MASH and cirrhosis. ^1,2^ Hepatic and peripheral insulin resistance are also defining features of T2D, along with fasting hyperglycemia, deficient pancreatic insulin secretion, and systemic changes in inflammatory profiles.^3,4^ Underlying drivers of both diseases are multifaceted, involving interplay between genetics, diet, exercise, and gene-environment interactions arising from various stressors (*e.g.,* infection, environmental exposure). ^3^ T2D is a heterogeneous disease with a spectrum of patient phenotypes, suggesting subgroups of patients may ultimately require different therapeutic interventions. ^3^ Translation of findings from preclinical animal models to therapeutic efficacy in humans remains poor ^5^, thus motivating new approaches to measure salient phenomena in patients, and more sophisticated humanized *in vitro* models. ^6^

Direct actions of insulin in healthy liver lead to inhibition of gluconeogenesis, activation of glycogen synthesis, and upregulation of lipogenic gene expression. ^4^ Insulin also acts indirectly on the liver by decreasing lipolysis in adipose tissue, resulting in reduction of circulating free fatty acids (FFA) and glycerol, further decreasing hepatic gluconeogenesis. ^4^ In T2D subjects, hepatic gluconeogenesis and FFA release from adipose tissue remain constant even in the presence of insulin, resulting in hyperglycemia and hepatocellular triglyceride accumulation. Chronically elevated levels of insulin and loss of receptor stimulation may enhance internalization/degradation, further altering acute metabolic activity and metabolic gene expression. This is compounded through indirect FFA and carbohydrate flux, influencing both acute activity and chronic expression of metabolic genes.^3,4,7^

Although insights into mechanistic underpinnings of insulin resistance in humans continue to emerge from well-crafted clinical studies ^8–10^, new experimental models are needed to parse the multi-variate contributions in different patient groups, and to better translate model predictions into therapeutic efficacy. The emergence of microphysiological systems (MPS), combining advances in 3D culture, microfabrication, and mesofluidics, has driven development of diverse approaches to extend functional performance of primary hepatocytes alone or in co-culture with non-parenchymal cells (NPCs), allowing *extended* and *mature* culture periods. Several of these MPS technologies have been deployed to model facets of insulin action, MASH, and T2D. ^11,12^ These models achieve a metabolically reactive disease state through modulation of glucose, free fatty acids, and insulin concentrations. However, among *in vitro* models, both the physiological and disease state is defined idiosyncratically, especially with respect to insulin concentrations. Insulin media concentrations, when reported at all, can range up to 1000 (or more)-fold greater than typical values in human plasma. ^13^

To address this, we parse the contributions of insulin and nutrient concentrations on the development and reversibility of insulin resistance in a primary human hepatocyte model. We implemented a liver MPS platform, that features microperfusion of cells cultured in a 3D scaffold, as this platform supports sustained human liver cell function under physiological conditions ^14^ and has been used for modeling human liver pharmacokinetics ^15^, metabolic diseases ^16,17^, and inflammatory conditions. ^18,19^ The utility of this platform for long term retention of select human liver functions, compared to other culture methods, was recently described by the FDA. ^20^ Of interest, the pharmacokinetic capability of this platform enables ready analysis of hepatocellular metabolic flux; insulin clearance, glucose production, and bile acid production, which are altered in T2D patients in heterogeneous ways across different patient populations. ^7^

Here, we establish a physiologically relevant metabolic baseline for hepatocytes *in vitro*. We then subsequently parsed the effects of hyperinsulinemia and nutrient overload, alone as well as in combination, to elicit and study the development, progression, and reversibility of insulin resistance during long-term culture.

## Methods

### Liver MPS Assembly and Operation During Culture

Two nearly identical (Liverchip™ (re-useable) and the disposable PhysioMimix™ (both from CNBio Innovations, Cambridge, UK)) versions of a polycarbonate micro-machined platform supporting 12 continuously perfused liver MPS-channels were used, according to already established methods (**Fig. S1**).^15,18^ The fluidic wells were primed by adding 1.6 mL priming media (1% BSA, 1% PenStrep in PBS) to each well continuously at 1 µL/s for 24 hours at 37°C. Priming media was aspirated to the level of the retaining ring immediately ahead of seeding. Following the seeding phase (*see below*), flow was maintained at 1 µL/s. Recirculating media were replaced every 48 hours by aspirating old media, adding 400µL new media for a 3-minute downward flow rinse step, followed by a full 1.6 mL media replacement.

### Cells and Media

The study was performed with primary human hepatocytes from a 50 y/o male donor (BioIVT; lot #SMC). Select experiments were repeated with cells from a 63 y/o male donor (BioIVT, lot #AQL). Cells were thawed in Cryopreserved Hepatocyte Recovery Media (CHRM, ThermoFisher), spun down at 100x*g* for 8 minutes, and resuspended in seeding medium; William’s E with 2 mM GlutaMax, 15 mM HEPES, 5% FBS, 1% P/S, 100 nM hydrocortisone, and glucose and insulin supplemented at levels corresponding to the proper experimental group. Typically, cells were seeded by adding 600,000 cells in 300 µL seeding medium to each well, allowing the cells to settle into the scaffold, and then adding an additional 1.3mL of seeding media (for total reactor volume of 1.6mL). Experiments optimizing clearance as a function of cell concentration were performed in triplicates using primary human hepatocytes (lot HU2098, ThermoFisher): 100,000; 200,000; 400,000; 600,000. Flow was initiated in a downward 1 µL/s mode for 8 hours to facilitate cell attachment to the scaffold and then reversed to an upward 1 µL/s flow for the remainder of the experiment. At 24 hours post-seeding, cells were switched to maintenance media; William’s E plus 6.25 µg/mL transferrin, 6.25 ng/mL selenium, 0.125% fatty acid-free BSA (18.8 uM), 2 mM GlutaMax, 15 mM HEPES, 100 nM hydrocortisone, 0.5% PenStep, and free fatty acids (FFA), glucose, and insulin, with concentrations adjusted to mimic various healthy and disease-inducing states. ^21^ Media states were designated as follows: “Condition 1” (200 pM insulin, 5 mM glucose, 20 µM FFA) “Condition 1 + FFA” (200 pM insulin, 5 mM glucose, 100 µM FFA), “Condition 1 + G + FFA” (200 pM insulin, 11 mM glucose, 100 µM FFA), and “Condition 2” (800 pM insulin, 11 mM glucose, 100 µM FFA). Baseline glucose of 5 mM corresponds to healthy fasting blood glucose levels and the elevated level of 11 mM corresponds to fasting blood glucose levels of diabetic patients. ^22^ FFAs were supplemented at a baseline of 20 µM of linoleic acid (normal maintenance condition for primary human hepatocytes) or 100 µM (with a physiologic ratio of 25% poly- unsaturated, 30% saturated, 45% mono-unsaturated e.g. 27.5 μM linoleic acid (18:2), 33 μM palmitic acid (16:0), and 49.5 μM oleic acid (18:1)) based on human plasma lipid composition. ^23^ Physionormic (200 pM) and high physiological (800 pM) insulin concentrations stem from well-established collected normal and T2D human sera values. ^24^

### Insulin Clearance

Insulin concentrations were measured by ELISA (Abcam, ab200011) in 50 µl media samples collected at 2, 4, 8, 24, and 48 hours after full media replacement. The clearance fraction was calculated by dividing insulin concentrations in collected samples by the initial concentration.

### Hepatic glucose production

Cells were first transitioned to a basal state by exchanging the medium to a formulation containing no glucose and no insulin and circulating for an hour. Media was then replaced with glucose-free media + 20 mM lactate and 1 mM pyruvate + insulin (0, 0.01, 0.1, 1, 10, or 100 nM as separate conditions). Cells were maintained in “glucose output” media for 24 hours followed by media collection for glucose concentration determination using the Amplex Red kit (Invitrogen, cat. no. A22189) or transcriptomics (see below).

### Cell Harvest for Molecular Assays

Endpoint analysis of cellular molecular properties (RT-qPCR, transcriptome sequencing, phospho-AKT, intracellular triglycerides, and metabolomics) was accomplished by lysing cells from the scaffolds at the end of each experiment. A unique lysis protocol was required for each type of assay. In cases where multiple analytes were measured in a single experiment, scaffolds were divided evenly between each assay to ensure at least *n*=2 technical replicates per assay.

### RT-qPCR

Scaffolds were rinsed in chilled DPBS and snap-frozen in liquid nitrogen. RNA was isolated using 1mL Trizol per scaffold. Scaffolds were sonicated for 30 seconds before adding 200µL chloroform and spun down at 18,000*xg* for 20 minutes. Supernatant was transferred and mixed with 500µL IPA for 10 minutes and spun at 18,000*xg* for 15 minutes. Pellets were rinsed twice with 75% EtOH and eluted into RNAse-free water. RNA was converted to cDNA using High-Capacity RNA-to-cDNA kit (Life Technologies, cat. no. 4387406) and qPCR was performed using TaqMan Fast Advanced Master Mix (cat. no. 4444556) with probes for *PCK1* (Hs00159918_m1), *G6PC* (Hs00609178_m1), and internal reference *18S* (Hs03928990_g1). Data were analyzed for fold change using the delta-Ct method^25^ and normalized to the 0 nM insulin group previously incubated in Condition 1 media.

### pAKT

Phosphorylation of AKT (AKT Serine/Threonine Kinase 1) was measured by starving cells for 30 minutes in insulin-free media and stimulating with insulin in basal media at doses ranging from 0nM to 100nM for 30 minutes. pAkt was measured using Milliplex MAP kit for total and phospho-AKT 1/2 (Millipore, cat. no. 48-618MAG) according to manufacturer instructions. After 30 minutes of insulin stimulation, cells were transferred immediately to kit lysis buffer for 30 minutes at 4°C with 1:100 protease inhibitor (Millipore, cat no. 535140), spun down at 3,000*xg* for 5 minutes, and collected for downstream quantification. Equivalent amounts of protein were determined and loaded in each assay using normalized BCA quantified lysate concentrations. For each biological replicate, the phospho/total Akt ratio was plotted and fit using a 3-parameter logarithmic curve in GraphPad Prism (version 8).

### Intracellular Triglycerides

Intracellular triglycerides were quantified using the Zen-Bio TG kit (Zen-Bio, cat. no. TG-1-NC) according to manufacturer instructions. Cells were lysed in 150 µL lysis buffer for 30 minutes at 4°C and frozen at -80°C. Upon thawing, 150 µL of wash buffer and 40 µL reagent B were added to scaffolds and incubated at 37°C for 2 hours. Lysates were diluted 1:2 in assay wash buffer followed by reagent A for detection of glycerol at 540 nm using a standard plate reader.

### Analysis of Bile Acids

MPS were disassembled and scaffolds with cells were extracted, rinsed once with 0.9% NaCl, and immediately placed in tubes with 1 mL ice-cold LCMS-grade isopropanol (Thermo Fisher) containing 100 nM d4-cholic acid (CDN isotopes) as an internal standard. Samples were vortexed for 10 minutes at 4°C and cleared by centrifugation at 8000 *x* g for 10 minutes at 4°C. 500 uL of supernatant was transferred to a new tube and evaporated under a stream of nitrogen. Dried extracts were resuspended in 50 µL acetonitrile/water (50/50 volume/volume) and 5 µL were injected into an ACUITY BEH C18 column (Waters, 3 x 100 mm x 1.7 μm). The HPLC system was a Dionex UltiMate 3000 HPLC coupled to a QExactive mass spectrometer equipped with an Ion Max source and a HESI II probe (Thermo Fisher Scientific, San Jose, CA). External mass calibration was performed using the standard calibration mixture every 7 days. Mobile Phase A was 0.1% formic acid in water and Mobile Phase B was 0.1% formic acid in acetonitrile. The column oven was set at 40°C and the autosampler was kept at 4°C. The flow rate was 0.6 mL min^-1^ and the gradient was as follows: 0-1.5 mins.: hold 2% B; 1.5-2.0 mins.: linear gradient 2- 35% B; 2.0-15.0 mins.: linear gradient 35-75% B; 15.0-15.1 mins.: linear gradient 75-100% B; 15.1-17.0 mins.: hold at 100% B; 17.0-17.1 mins.: linear-gradient 100-2% B; 17.1-19.0 mins.: hold at 2% B. Data were acquired in full-scan, negative mode, with a scan range of *m/z* = 200-600. The resolution was 70,000, with an AGC target of 1x10^6^, and IT at 80 ms. Relative quantitation of bile acids were performed using TraceFinder™ 4.1 (Thermo Fisher Scientific) using a 5 ppm mass tolerance and referencing an in-house library of chemical standards. Raw peak areas were normalized to d4-cholic acid.

### Transcriptome Analyses

Total RNA from hepatocyte scaffolds were extracted as previously detailed (*see RT-qPCR*). RNA purity and concentration were determined using a Nanodrop One (Thermo-Fischer Scientific). The preparation of RNA library and whole transcriptome sequencing was performed by EuroFins. The method for differentially expressed gene (DEG) determination was edgeR using the open-source R/BioConductor software.^26^ Gene set enrichment analysis (GSEA) and clusterProfiler (Gene Ontology: Biological Process) were used to detect pathway-level differences among sample sets. ^27,28^ The effects of separate RNA isolation dates were treated as covariates and batch-corrected using the ComBat normalization method. ^29^

### Statistical Analysis

All data shown are mean ± SD. All statistical comparisons were performed using two-way ANOVA in GraphPad Prism version 8. If the interaction term was not significant, *p* values were reported for the column factor (typically experimental condition, # *p* < 0.05, ## p < 0.01, ### *p* < 0.001) and row factor (either dose-response or time course, $ *p* < 0.05, $$ p < 0.01, $$$ p < 0.001). If the interaction term was significant, post-hoc multiple comparison analyses were run using either Tukey test (3 or more groups) or Sidak test (2 groups). For dose-response curves with 3 or more groups, Tukey post-hoc test compared the main effect between experimental conditions. For time courses or dose-response curves with only 2 experimental groups, a post-hoc test compared the experimental condition effect within each time point or dose. For all post-hoc tests, *p* values were reported for the post-hoc test only (* p < 0.05, ** p < 0.01, *** p < 0.001, **** p < 0.0001). For insulin dose response, a nonlinear regression 3 parameter fit ([Agonist] vs. dose-response) was performed to determine IC50 values. For metabolomics data, the discovery of significant metabolites was performed using multiple t-tests in GraphPad Prism of normalized expression values corrected for multiple comparisons using an FDR threshold of 10%.

## Results

### Hyperinsulinemia induced insulin resistance in vitro

Primary human hepatocytes were cultured for 7-15 days in the liver MPS in either baseline media (Condition 1; 200 pM insulin, physiological nutrients) or in Condition 1 + hi-insulin media (Condition 1 + Ins; 800 pM insulin, physiological nutrients), reflecting the reported human portal vein insulin concentrations in healthy and T2D patients ^30,31^. Insulin clearance and albumin production rates were monitored between regular 48-hour feeding periods, and hepatocyte glucose production (HGP) was evaluated over 24 hours in glucose-free media with insulin dose-response (0-100 nM) assayed from days 7-8 and 14-15 of the experiment (*see Methods*). Supporting the notion that hepatocytes cultured in our device are molecularly ‘mature’ and in agreement with previous findings ^16,17^, hepatocytes exhibit stable albumin secretion rates over 15 days of culture following an initial transient increase, with no differences in magnitude among baseline and high insulin media conditions or between LiverChip MPS models (**Fig. S2**).

Insulin clearance rates as a fraction of total insulin present were comparable in both 200 pM and 800 pM insulin-treated hepatocytes for the first week of culture (**Fig. 1A**), suggesting a first-order clearance rate consistent with the reported 190 pM Kd for high-affinity insulin binding to the insulin receptor. ^32^ Reactors seeded at lower cell numbers (400,000; 200,000; or 100,000) showed roughly proportional reductions in clearance compared to the standard seeding density of 600,000 cells per reactor (**Fig. S3**). The per-cell clearance rate at the nominal 200 pM and 800 pM are between 350-1500 molecules/min, well within a feasible range for trafficking of the insulin receptor. ^33^ Cells maintained in baseline conditions retained their ability to clear insulin at close to initial rates throughout the 14-day culture period, with a modest decline from 80% to 77% (**Fig. 1A**). In contrast, hepatocytes maintained in 800 pM insulin lost almost half their initial clearance rate by day 14, starting on day 9 (**Fig. 1A**).

**Figure 1.**
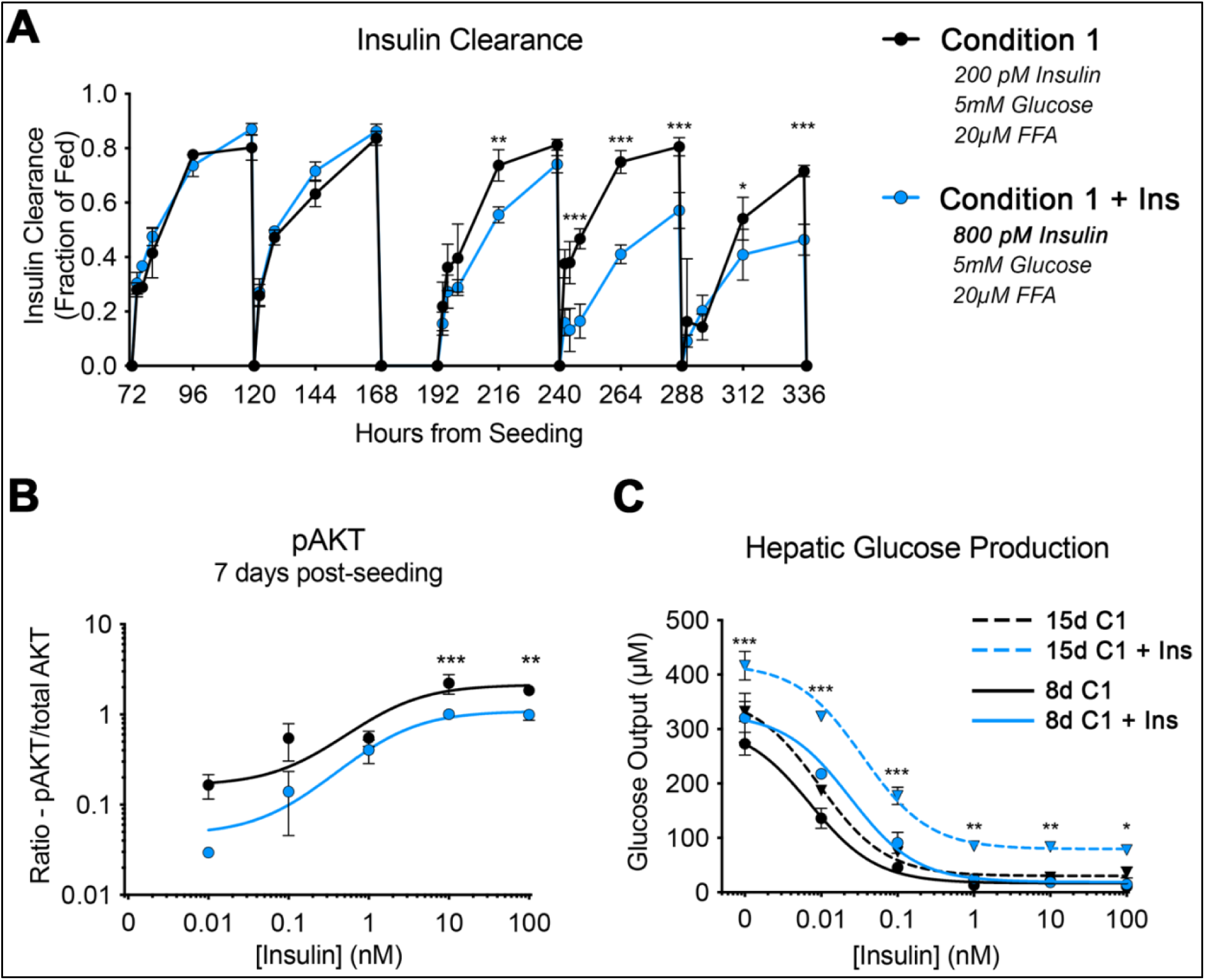
Hyperinsulinemia alone induces insulin resistance in 3D perfused MPS culture. (A) Cells in hyperinsulinemic conditions (Condition 1 + Ins; 800 pM insulin) gradually lose the ability to clear insulin over 15 days in culture compared to (Condition 1; 200 pM insulin). Two-way ANOVA interaction term p < 0.001, and asterisks indicate significance in post-hoc Sidak test between 800 and 200 pM maintenance media (* p<0.05, ** p<0.01, *** p<0.001). (B). Cells in hyperinsulinemia condition have reduced responsiveness to insulin stimulation, as assessed by activation of the downstream signaling target AKT, using phosphorylation as a surrogate for activation. Two-way ANOVA was significant for maintenance media term (p<0.0001) and post-hoc Sidak’s test between 800 and 200 pM maintenance media (*** p<0.001, ** p<0.01). (C) Cells maintained in hyperinsulinemia show decreased suppression of hepatic glucose output upon 24-hour insulin stimulation. At day 8, two-way ANOVA was significant for insulin dose response ($$$ p<0.001 for row factor) and maintenance media (## p<0.01 for column factor). At day 15, glucose output was significantly higher in 800 pM insulin maintenance media than 200 pM insulin maintenance media at all insulin dose responses (interaction term p<0.01, post-hoc Sidak *** p<0.001, ** p<0.01, * p<0.05).

The decrease in clearance was paralleled by a corresponding reduced ability of cells in the hyper-insulinemic state to initiate downstream signaling when stimulated with insulin, as assessed by phosphorylation of the downstream insulin receptor target, AKT (**Fig 1B**). Similarly, cells maintained in 800 pM insulin displayed increased basal gluconeogenesis and reduced insulin sensitivity by days 8 and 15, as measured by insulin-induced suppression of HGP, compared to cells maintained in 200 pM insulin. This impairment was observable at only the lowest insulin doses used in the HGP assay on day 8, but widely observed in all insulin doses by day 15 in hepatocytes maintained in high insulinemic conditions (**Fig. 1C**). Supporting the replicability of these assays, congruent results are obtained from experiments performed using hepatocytes obtained from a separate donor (**Fig. S4A, B**) and in slightly different bioreactors (**Fig. S2C**). In sum, hepatocytes cultured in hyperinsulinemic conditions within our *in vitro* MPS model develop reduced insulin clearance, which alters downstream AKT signaling cascades, and impairs HGP.

### Hyperglycemia- and hyperlipidemia-induced insulin resistance

Having established that hyperinsulinemia alone induced several features of the insulin-resistant phenotype in our *in vitro* platform, we then assessed hepatocellular responsiveness following exposure to other common agonists of metabolic disease; FFAs, Glucose, alone or in combination with hyperinsulinemic conditions. Condition 1 (200 pM insulin, 5 mM glucose, 20 µM FFA), Condition 1 + FFA (100µM), Condition 1 + G (glucose: 10mM) + FFA (100µM), and Condition 2 (800 pM insulin, 10 mM glucose, 100 µM FFA) media formulations were used to assay hepatic insulin clearance and HGP through 19 days of culture. By day 19, FFA stimulation alone (Condition 1 + FFA) leads to the highest IC50 (.052) value of the three metabolically stimulated conditions (**Fig. 2B**). However, only hepatocytes treated with Condition 2 display a significantly heightened glucose production across all doses and all time points (12d: IC50_Condition 1_ = 0.0076 vs IC50_Condition 2_ = 0.027, 19d: IC50_Condition 1_ = 0.0087 vs IC50_Condition 2_ = 0.028) (**Fig. 2A**). In congruence with this data, RT-qPCR of two gluconeogenic genes, *PCK1* and *G6PC*, ^34^ revealed hampered transcriptional repression in hepatocytes maintained in either Condition 1 + G + FFA or Condition 2 compared to cells in Condition 1 following a 24-hour incubation with 0.1 and 1nM Insulin (**Fig. 2B**). Thus, even in the absence of hyperinsulinemia, high glucose/FFA results in a protein and transcriptional impairment in the hepatocellular response to insulin. Conversely, insulin clearance by hepatocytes maintained in the other nutrient agonists is reduced only if hyperinsulinemia is concomitantly present (12d: IC50_Condition 1_ = 0.0087 vs IC50_Condition 2_ = 0.028, 19d: IC50_Condition 1_ = 0.012 vs IC50_Condition 2_ = 0.023)(**Fig. 2C**), suggesting high insulin as the primary driver of metabolic impairment within our culture model.

**Figure 2.**
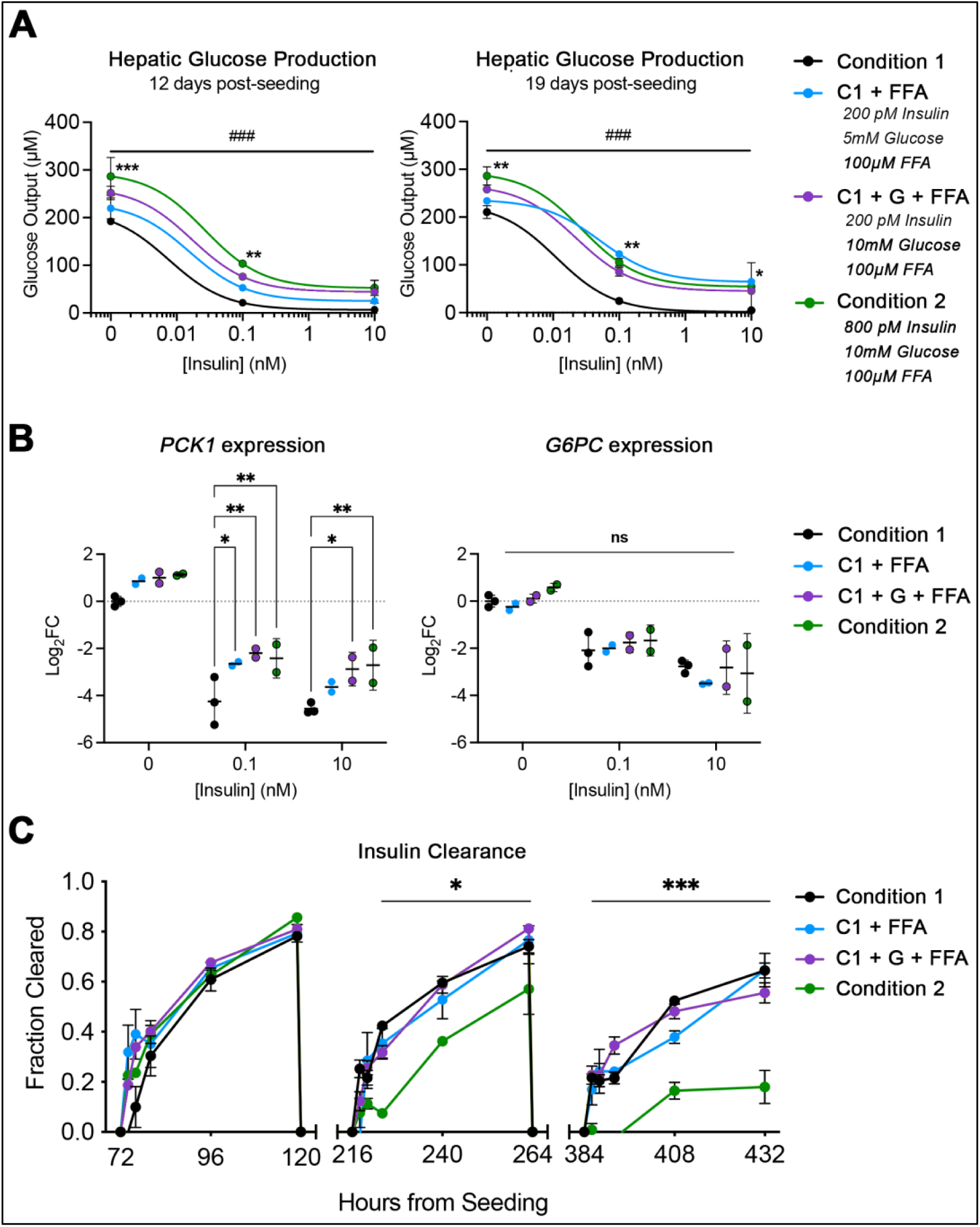
Nutrient overload induces insulin resistance. (A) Glucose output is significantly increased in both Condition 1 + FFA and Condition 2 over the baseline condition on days 12 and day 19. ### p<0.001 for the dose variable, and * p<0.05, ** p<0.01, *** p<0.001 via two-way ANOVA. (B) Condition 2 has impaired *PCK1* responsiveness to insulin at .1 and 10nM comparied with Condition 1. G6PC expression is similar across all doses and sample-sets. * p<0.05, ** p<0.01 *** p<0.001 via two-way ANOVA. (C) Insulin clearance is significantly different between Condition 2 and Condition 1 at 9 and 16 days incubation. Two-way ANOVA; * p<0.05, *** p<0.001. *n*=4; all samples.

### Individual high-nutrient conditions further exacerbate hyperinsulinemia-induced insulin resistance

Leveraging these insights, we individually assayed which metabolic agonist, glucose or FFA, further influenced the various insulin resistance metrics in addition to hyperinsulinemic conditions. Insulin clearance was evaluated during each feeding period, and HGP was monitored between days 7-8 and days 14-15. Consistent with **Figs. 1 & 2**, adding either hi-glucose (Condition 1 + Ins + G), or both high glucose/FFA to the high insulin media (Condition 2) does not reduce insulin clearance compared to high insulin alone (**Fig. 3A**). Interestingly however, it is the addition of both high glucose *and* high FFA that impairs insulin sensitivity the most to transcriptional and protein-level HGP suppression beyond that for hyperinsulinemia alone (8d / 15d: IC50 _Condition 1+Ins_ = 0.025 / 0.090 vs IC50 _Condition 1+Ins+G_ = 0.041 / 0.065 vs IC50_Condition 2_ = 0.051 / 0.065)(**Fig 3B, C**). Thus, hepatocytes in our model phenotypically recapitulate insulin resistance most aptly through joint incubation with high insulin, glucose, and FFAs; *Condition 2*.

**Figure 3.**
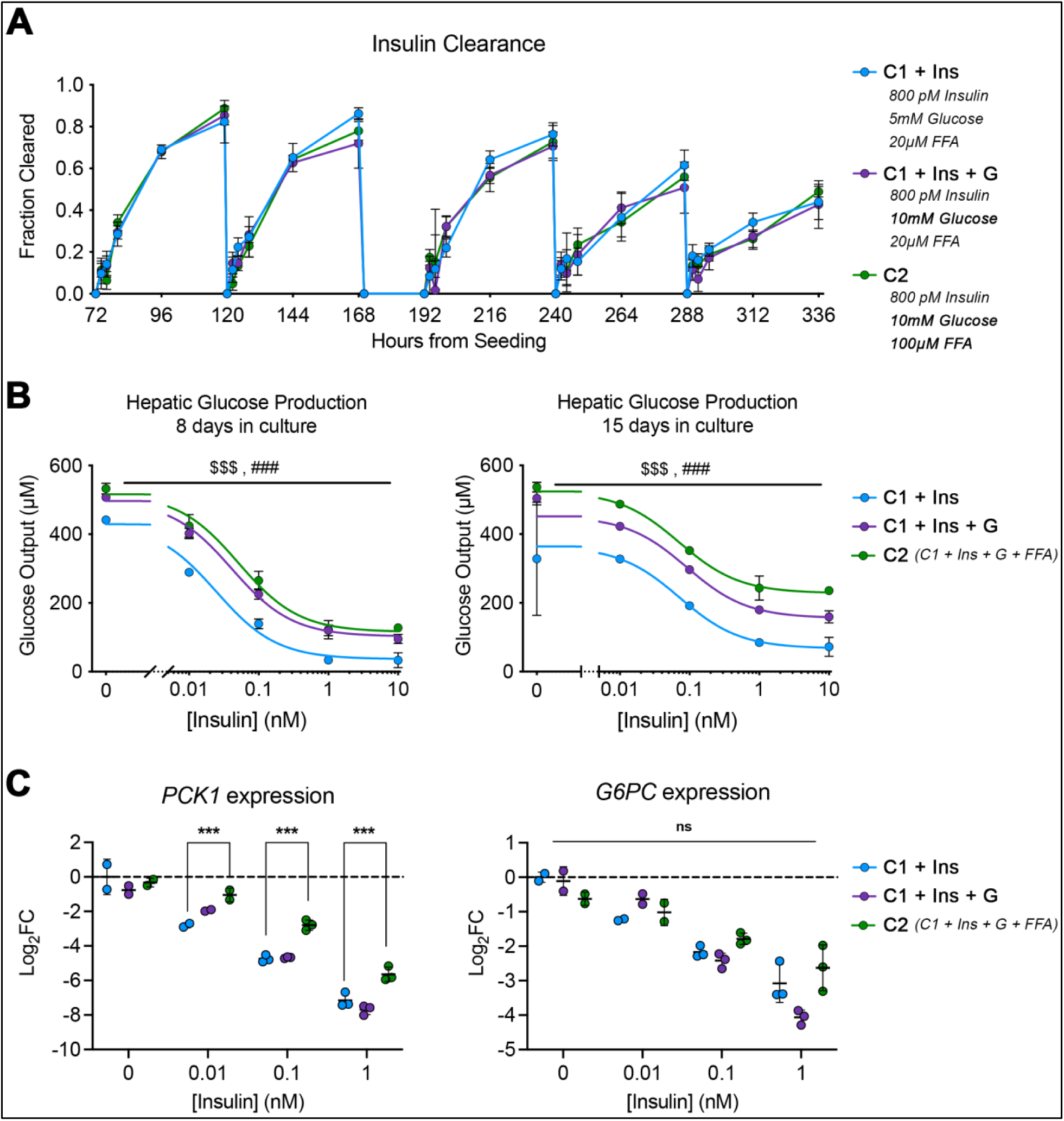
Nutrient overload induces insulin resistance. (A) Insulin clearance decreases over time in culture in all groups, but there is no additive effect of high glucose or high FFA on insulin clearance. (B) Both Condition 1 + Ins + G, and Condition 2 have significantly increased glucose output over Condition 1 + Ins hepatocytes at days 8 and 15 via two-way ANOVA ($$$ p<0.001 for dose variable and ### p<0.001 for maintenance media variable). (C) All groups show decreased gluconeogenic gene expression with increasing insulin dose, however, Condition 2 cells have significantly higher expression of *PCK1* after incubation with 0.01, 0.1, and 1 nM insulin than Condition 1 + Ins alone (p<0.001). No difference in *G6PC* expression was noticed amongst groups; two-way ANOVA.

### Transcriptome analysis of hepatocellular insulin resistance in vitro

Having determined that Condition 2 induces core aspects of the insulin resistance phenotype, RNA sequencing analysis of hepatocytes incubated in Condition 1 and Condition 2 media formulations identified 914 differentially expressed genes (DEGs; *p*adj < 0.05), with condition-specific separation by principal component analysis (PCA; **Fig. 4A**). Of the 914 DEGs, 38 and 27 had a Log_2_FoldChange > 1 or < -1, respectively (**Fig. 4B**). Significant downregulation of metabolic mediators; *FOXQ1*, *IGFBP1*, *SERPINE1*, and *ADM*, and upregulation of pro-inflammatory cues; GFAP, BMF, and FGF1, indicate a reactive hepatocellular cell state following two weeks in Condition 2 media (**Fig. 4B**). Expanding this analysis, pre-ranked gene set enrichment analysis suggests upregulated gene contributions in the interferon-alpha response (**Fig. 4C, D**). Most prominently, there is significant basal upregulation of chemokine *CXCL10*, and stress/metabolic mediator *TXNIP*. Consistent with historical findings of NF-kB mediation of insulin sensitivity, ^35,36^ Condition 2 hepatocytes in our system have a noted downregulation in NF-kB target genes (**Fig. 4C, E**). Condition 2 hepatocytes also show expected significant metabolic dysfunction, as shown by transcriptional alterations of main CYP enzymes; *CYP3A4*, *CYP2A6*, and *CYP2A7* (**Fig. 4F**), and concomitantly increased transport/synthesis of bile acids; *ABCB4* (MDR3), *AKR1D1*, and *CYP7A1* (**FIG. 4G**). In sum, hepatocytes exposed to insulin/nutrient overload are pro-inflammatory and have perturbed metabolic signaling cascades, likely predisposing them to the observed attenuated response to insulin (**Figs. 1-3**).

**Figure 4.**
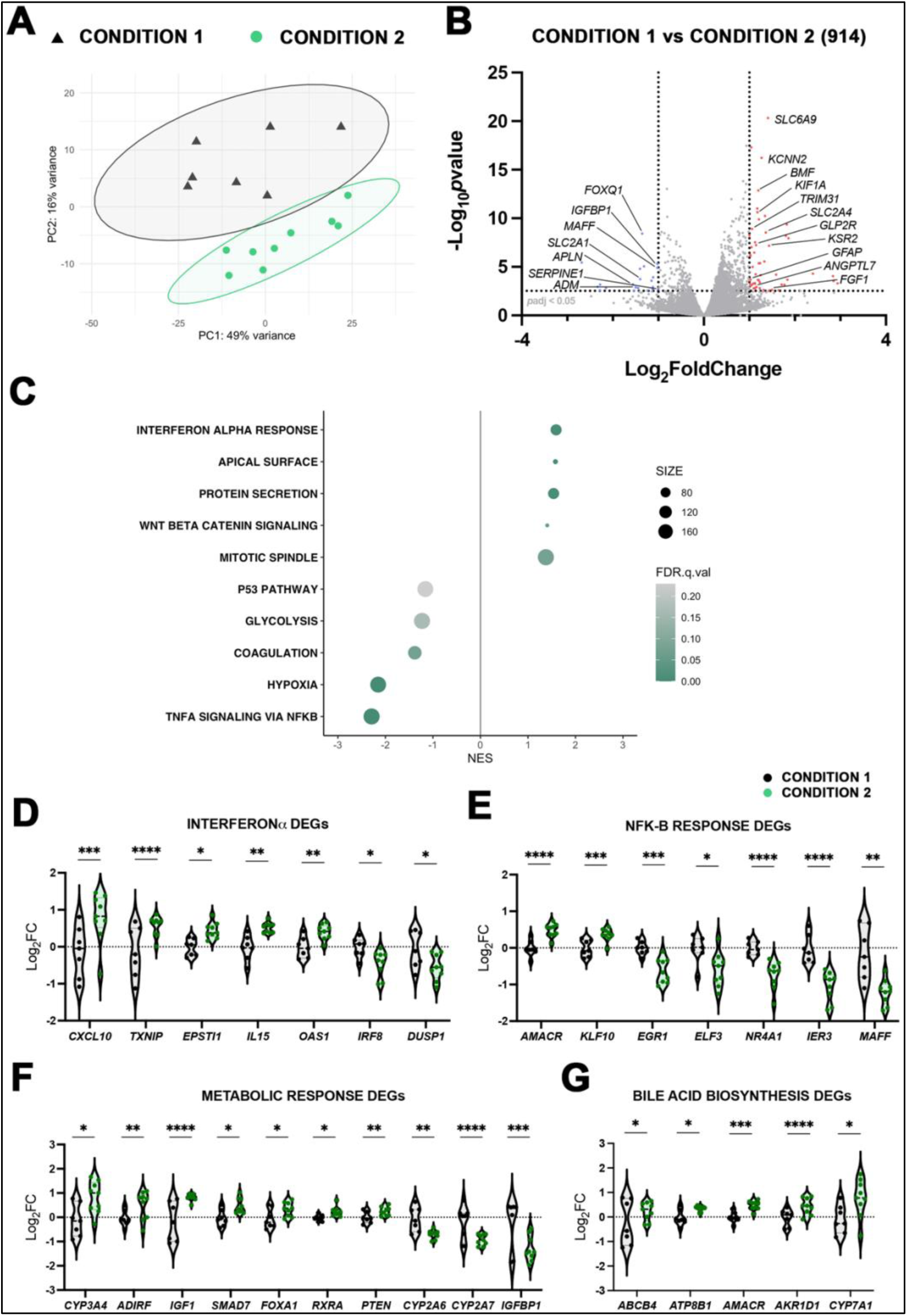
Activation of transcriptional signaling pathways in insulin-resistant primary human hepatocytes. (A) Prinicipal component analysis (PCA) separates samples by exposure to 14 days culture in Condition 1 (*n* = 7) and Condition 2 (*n* = 9). (B) Volcano plot of log_2_ fold change and p values between Condition 1 and Condition 2 transcripts. Red dots and blue dots indicate transcripts that had both a padjusted < 0.05 and log_2_ fold change > 1 or < -1, respectively. Gray dots are transcripts that remain relatively unaffected between samples. (C) Hallmark gene pathways enriched in Condition 1 and Condition 2 were identified by gene set enrichment analysis. (D-G) Select differentially expressed genes (DEGs) log_2_ fold changes were plotted in pathways regulating Interferon-α, NF-kB, metabolic, and bile acid response. * *p*adj<0.05, ** *p*adj<0.01 *** *p*adj<0.001, **** *p*adj<0.0001 acquired from RNA sequencing data.

To further parse this hypothesis, that our *diseased* hepatocytes are metabolically poised to an impaired insulin response, we exposed Condition 1 and Condition 2 to 1nM of insulin for 24 hours after two weeks in MPS culture and performed RNA sequencing on their cell lysates (**Fig. 5A, B**). Both hepatocyte cultures separated via PCA following insulin treatment (**Fig. S5A, B**). Surprisingly, both had similar levels of DEGs, 1060 and 1079 in Condition 1 and Condition 2 after insulin treatment, respectively. To identify the genes that are differentially insulin-responsive between each condition, we plotted the DEGs (*p*adj in either insulin-stimulated Condition 1 or Condition 2 cells) of both groups according to their response similar index (RSI)(**Fig. 5C**), which colors DEGs by their signed probability of joint differential expression, where values near 1 indicate a high probability of concordance, and near -1 are likely discordant. ^37^ DEGs were additionally separated by magnitude (|Log_2_FC_Condition 1 vs Condition 1 + Insulin 1nM_ - Log2FC_Condition 2 vs Condition 2 + Insulin 1nM_|) (**Fig. S5C**) to produce a focused list of 422 DEGs that are differentially concordant (RSI > 0, Mag > 0.25) between insulin-stimulated Condition 1 and Condition 2 hepatocytes (Supplemental Data). Interestingly, Gene Ontology (GO) enrichment of the focused DEGs identified *bile acid and bile salt transport* as the most differentially regulated biological process dividing the Condition 1 and Condition 2 hepatocellular response to insulin (**Fig. 5D**). Within this pathway, major bile acid biosynthesis/transport genes are differentially upregulated between our two groups following insulin treatment; *CYP7A1*, *ABCC4* (MRP4), *SLC51A* (OSTα), and *ABCB11* (BSEP) (**Fig. 5E**), aptly modeling the strong correlation found between elevated cholestatic markers and patients with insulin resistance. ^38,39^ Immune system mediators *AXL*, *GAS6*, and downstream *AKT3* were also among our list of concordant DEGs (**Fig. 5F**), suggesting a possible role for AXL-GAS6 in mediating insulin receptor trafficking. ^40^

**Figure 5.**
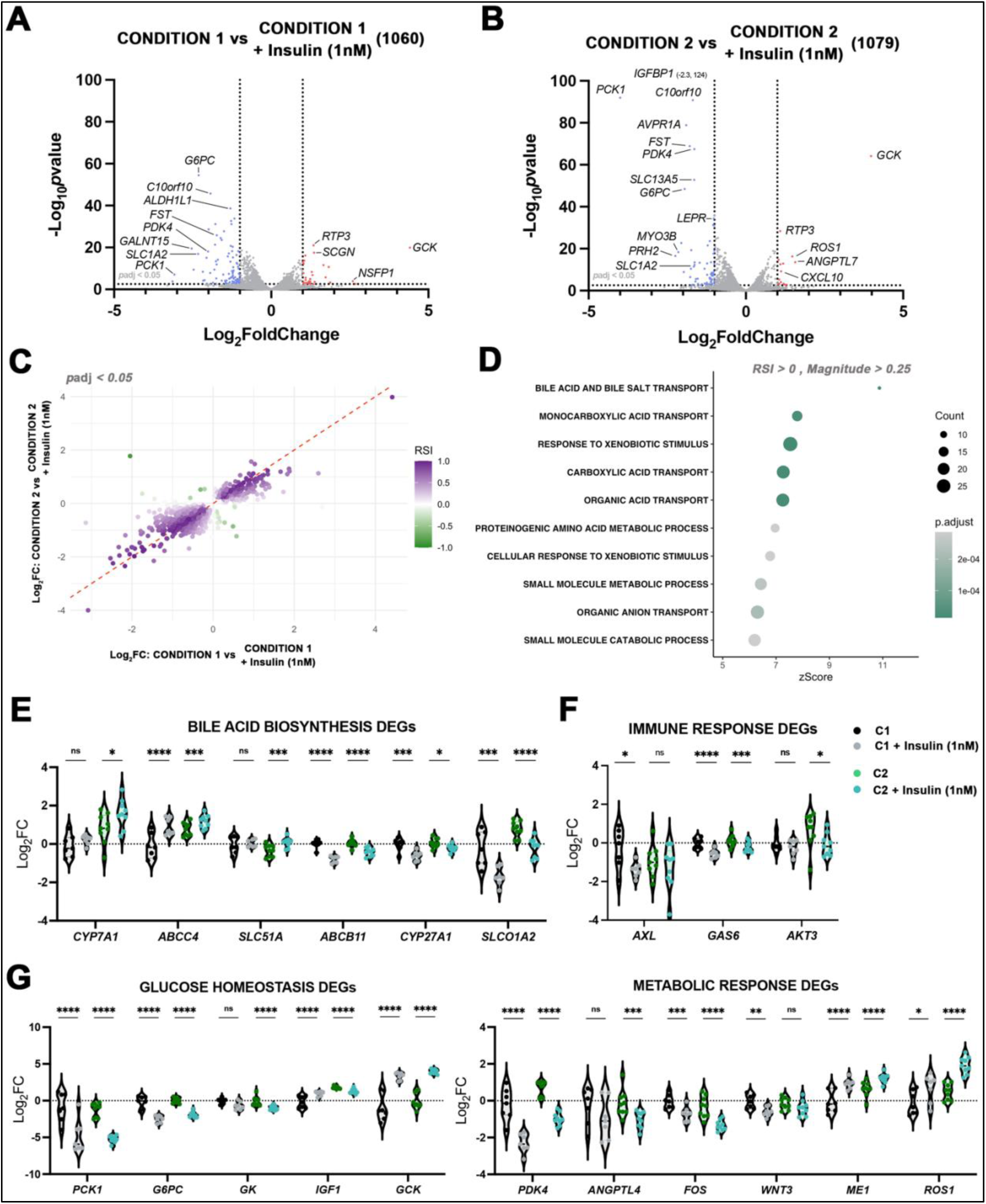
Transcriptionally impaired insulin sensitivity in hyperinsulinemic primary human hepatocytes. (A, B) Volcano plot of log_2_ fold change and *p* values between (*n* = 7) and Condition 2 (*n* = 9) transcripts following exposure to 1nm Insulin (Condition 1 +1nM Insulin [*n* = 7] and Condition 2 +1nM Insulin [*n* = 9]). Red dots and blue dots indicate transcripts that had both a padjusted < 0.05 and log_2_ fold change > 1 or < -1, respectively. Gray dots are transcripts that remain relatively unaffected between samples following insulin exposure. (C) Response similarity index outlining transcriptional concordance (values near 1, purple) and discordance (values near -1, green) between differentially expressed genes (DEGs) in Condition 1 and Condition 2 following incubation with 1nM insulin. (D) Concordant DEGs were enriched in multiple metabolic pathways following biological process Gene Ontology Enrichment. (E-G) Select differentially expressed concordant genes (DEGs with RSI > 0, magnitude > 0.25) log_2_ fold changes were plotted in pathways regulating bile acid, immune, glucose homeostasis, and metabolic response. * *p*adj<0.05, ** *p*adj<0.01 *** *p*adj<0.001, **** *p*adj<0.0001 acquired from RNA sequencing data.

Of importance, and within our differential concordant list of DEGs, *PCK1* and *G6PC* both had a shunted transcriptional response to insulin in Condition 2 hepatocytes (**Fig. 5G**; left), amongst other key glucose homeostasis genes. Subsequent metabolic mediators; *PDK4*, and *ROS1*, also had a significantly impaired response to insulin in Condition 2 hepatocytes compared to those in Condition 1 (**Fig. 5G**; right). In sum, these data outline a hepatocellular state modeling several aspects of insulin resistance and serve as a starting point for any anticholestatic or antidiabetic therapeutics considered in this approach.

### Elevated bile acids within insulin-resistant MPS hepatocytes

Corroborating these transcriptomic findings, we performed metabolomic analysis of hepatocytes in Condition 1, Condition 1 + Ins or Condition 2 at both an early (day 12) and late (day 19) timepoint, and identified a significant increase in taurine- and glycine-conjugated cholic acid in only Condition 2 treated hepatocytes compared to baseline after 9 days in our biorector (**Fig. 6A**). This phenotype tapered only for glychocholic acid after 18 days in culture, however (**Fig. 6B**). There were no significant differences in chenodeoxycholic acid or its conjugated forms (data not shown). This phenotype validates the observed increase in enzymes mediating transport/production of bile acids (**Fig. 4G** & **Fig. 5D, E**), suggesting a role for cholesterol catabolism in response to hyperinsulinemic/nutrient overload. ^41^

**Figure 6.**
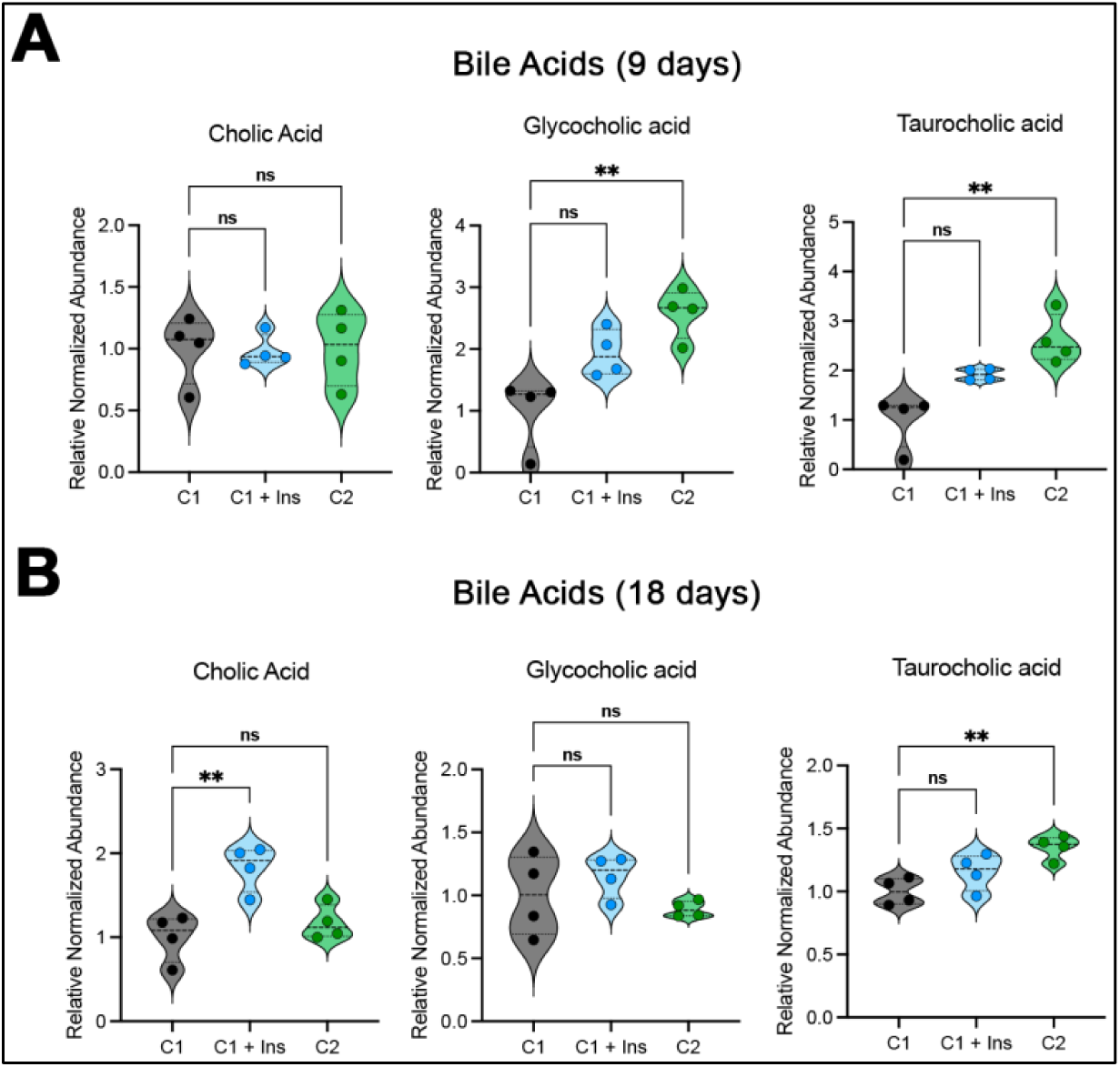
Insulin-resistant phenotype leads to bile acid metabolic dysregulation. (A) Intracellular abundance of primary conjugated bile acids is significantly increased at day 9 in both Condition 1 + Ins and Condition 2 vs. Condition 1 (Tukey’s post-hoc test, ** p<0.01, * p<0.05). (B) After 18 days in culture, taurocholic acid is still significantly elevated in Condition 2 over the Condition 1 stimulated hepatocytes (Tukey’s post-hoc test, * p<0.05).

### Incomplete reversal of hepatocyte insulin responsiveness upon return to Condition 1 media

Finally, to assess if insulin-resistant hepatocytes were capable of returning to baseline insulin sensitivity, hepatocytes were cultured in either Condition 1, Condition 1 + Ins, or Condition 2 media for 12 days, followed by a 50% supplement in Condition 1 (*recovery media*) for days 12-19. Cells cultured in the recovery condition for days 12-19 displayed HGP (**Fig. 7A**) and insulin clearance (**Fig. 7B)** that were nearly identical to cells maintained in Condition 1 media for the entirety of the 19-day experiment, regardless of which diabetes-mimicking media in which they were initially cultured. Conversely, hepatocyte intracellular triglycerides are persistently elevated in the Condition 2 and Condition 1 + Ins, even after supplementation with 12 days of *recovery media*, indicating insulin resistance may not be entirely reversible within our system, and the need for longer time points, or additional treatment options (**Fig. 7C**).

**Figure 7.**
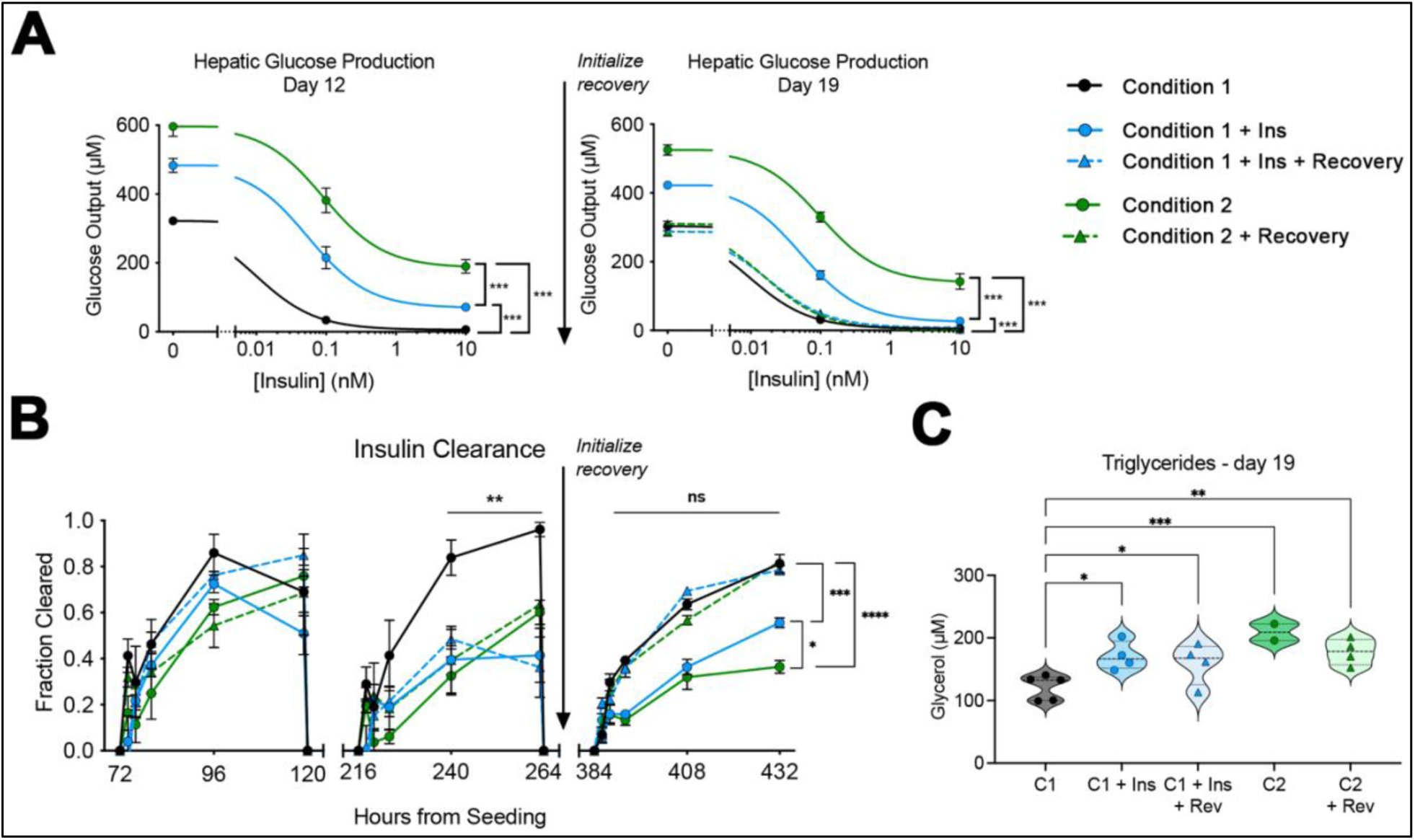
Partial recovery of insulin sensitivity after return to Condition 1 media. (A) Hepatic glucose output is significantly different between all groups at day 12 (*** p<0.001 Tukey test main column effect). At day 19, cells returned to Condition 1 from days 12-19 show no significant difference in glucose output, while cells maintained in high insulin (Condition 1 + Ins) or high insulin with nutrient overload (Condition 2) are still significantly different than all other groups and each other (*** p<0.001 Tukey test main column effect). (B) Insulin clearance is diminished in Condition 1 + Ins and Condition 2 by day 12 (hour 240-264) in culture (** p<0.01, Condition 1 vs all other groups). Upon return to healthy conditions, the recovery groups clear insulin similar to the healthy control (no significance; *ns*), while the chronic high insulin (Condition 1 + Ins) and Condition 2 hepatocytes continue to exhibit decreased insulin clearance by 18 days. (C) Intracellular triglyceride content at day 19 is significantly higher than the control in cells exposed at any time to Condition 2 media, regardless of whether the cells were returned to healthy media. * p<0.05, ** p<0.01, *** p<0.001 via two-way ANOVA.

## Discussion

Here, we used an established liver MPS model^15–20^ to parse the contributions of hyperinsulinemia and elevated nutrients on the development of the insulin resistance phenotype in primary human hepatocytes. Of note, primary human hepatocytes cultured in a physiological medium (Condition 1; 200 pM insulin, 5.5 mM glucose, 20 uM FFA) remain insulin sensitive over almost 3 weeks in culture, as demonstrated by insulin clearance, dose-dependent Akt phosphorylation, and dose-dependent suppression of HGP by insulin (**Figs. 1, 2, 6**). Second, a modest increase in insulin at the time of medium change, from 200 pM to 800 pM, induces an insulin-resistant phenotype over 7-10 days in culture (**Figs. 1, 4**), jointly validated with an additional donor set (**Fig. S4**). Hepatocytes cultured in hyperinsulinemic conditions have exacerbated phenotypes in the presence of hyperglycemia and elevated FFAs (**Figs. 2, 3**). Further, we identify the transcriptional changes in our *diseased* hepatocytes that poise these cells to an impaired insulin response (**Figs. 4, 5**), with protein-level concordance in secondary bile acid signaling (**Fig. 6**). Finally, the insulin-resistant model shows partial reversibility when cells are restored to baseline medium conditions (**Fig 4**). The results from such a succinct array of analyses indicate that the insulin-resistant hepatocytes within our established bioreactor fundamentally model multiple aspects of what is seen in the human insulin-resistant condition (**Fig. 8**; schematic).

**Figure 8.**
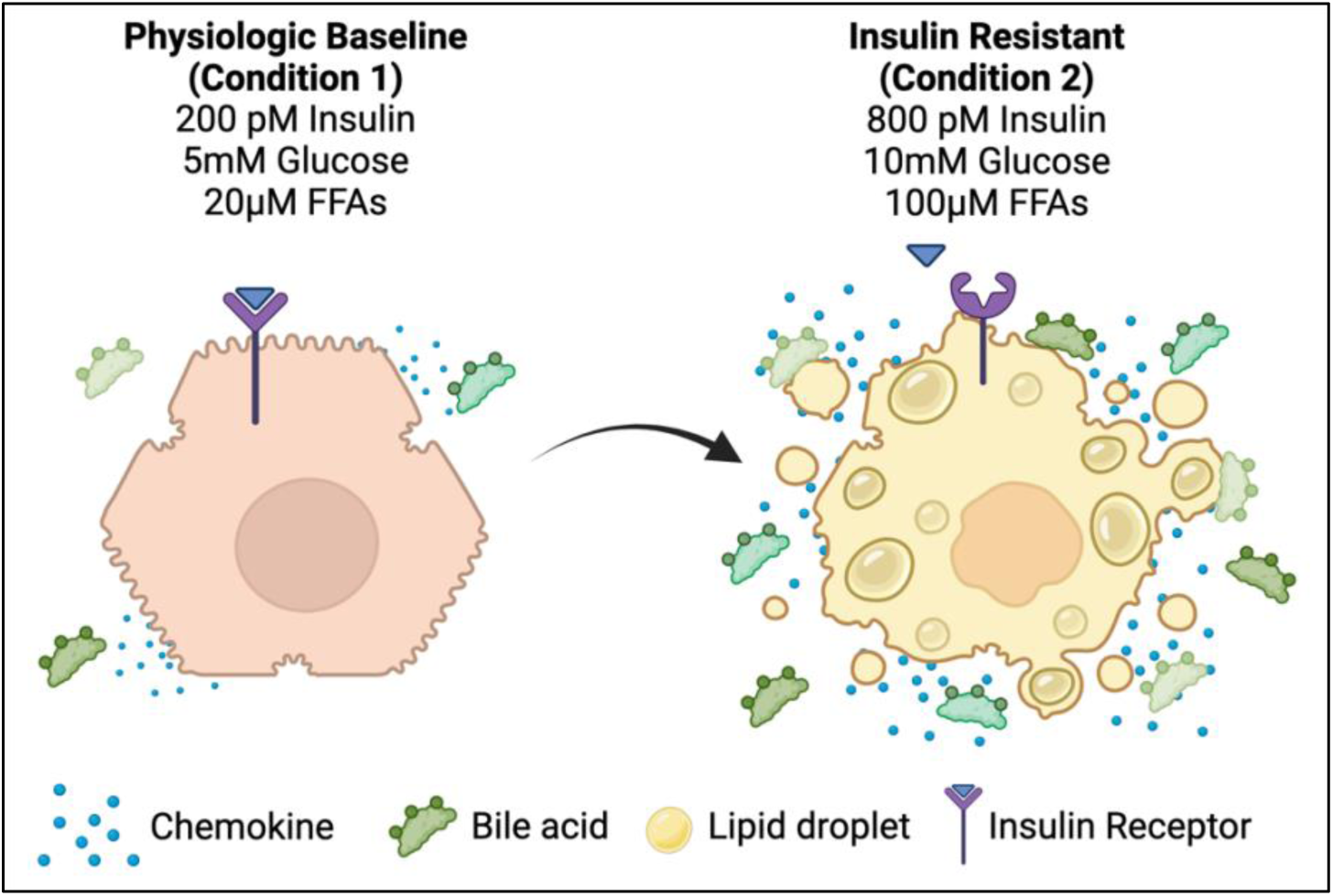
Schematic depiction of the physiological baseline versus insulin-resistant hepatocytes in our MPS model. On the left is a hepatocyte treated with physiological levels of insulin, glucose, and FFAs, mirrored by an insulin-resistant Condition 2 incubated hepatocyte on the right. Insulin-resistant hepatocytes are pro-inflammatory, contain increased intracellular triglycerides, and have elevated bile acid expression. This figure was made with BioRender.com under a purchased license agreement.

Our physiologically relevant healthy and diseased media formulation, which to the best of our knowledge, is the first instance of its implementation to *in vitro* primary human hepatocytes in an MPS, is significantly different and a stark improvement in comparison with other *in vitro* studies. Insulin concentrations of 200 pM for healthy and 800 pM for disease approximate portal vein values for these states integrated over fasting/fed excursions *in vivo* in healthy and diseased states. ^30,31^ Corroborating this choice, by elevating insulin concentrations *alone* to the values expected in T2D subjects, we recapitulated several features of the hepatic insulin-resistant phenotype (**Fig. 1**). The finding of reduced clearance in our models’ *disease* state is in accordance with multiple human studies reporting a decrease in insulin clearance in T2D subjects and isolated cells from T2D livers. ^42,43^ In comparison with reported *in vitro* hepatic disease models, most use supraphysiological insulin for both healthy- and disease states, ^44,45^ pair a physiological healthy value of nutrients (glucose/FFA) with supraphysiological for disease ^45,46^ or do not report insulin/nutrient values at all. ^17^ Further, clearance is not typically reported, perhaps because cell concentrations are either much lower, ^45,47^ higher, ^44^ or certain *in vitro* models lack pharmacokinetic capability. ^15,48^ As such, the choice of these physiologically reflective values of insulin, combined with the choice of our bioreactor, allows for reproducible measurements of clearance and should provide a baseline for assaying insulin sensitivity across all *in vitro* models. A more comprehensive picture of insulin, nutrient, and steroid values will likely emerge as studies standardize protocols for metabolic exposure including the complex interactions with serum proteins.

While elevation of insulin alone was enough to induce aspects of the insulin-resistant phenotype, the elevation of glucose or FFAs alone, or in combination, did markedly induce metabolic changes in the hepatocellular phenotype (**Fig. 3**), and as such are probably more reflective of a MASH or T2D condition. ^4^ By investigating combinations of these three metabolic agonists in physiologically normal and diseased levels, we uncover immunologically reactive hepatocytes (**Fig. 4. C-E**), that are significantly impaired in their response to insulin (**Fig. 5**). These transcriptomic signatures share strong similarities across other models and human datasets, ^37^ and are reflective of an early insulin resistant or MASH model; hampered response to insulin, but not as immunologically reactive as late-stage. ^49,50^ Perhaps longer hepatic incubation times in the Condition 2 media or inclusion of non-parenchymal cells would elicit such a response.

Additionally, by analyzing RNA-seq data using focused enrichment analysis, we identified bile-acid synthesis/transport as significantly upregulated before and after insulin treatment in our *diseased* hepatocytes. This phenotype likely comes secondary to a compensatory production of intracellular glycogen and nutrient overload within our system, with downstream catabolism of intracellular triglycerides. ^41^ Nonetheless, further therapeutic avenues should assess anticholestatic agonists, well known for their ability to induce bile acid secretion, reduce lipid content, and likely improve insulin sensitivity. ^51^ These data further posit this model as one that reflects multiple aspects of hepatic insulin-resistance, but warrant multiple follow-up studies investigating the complex regulatory effects of metabolic stress in driving aberrant insulin signaling and immunological shifts.

There are a few notable limitations of our *in vitro* model of human hepatic insulin resistance. First, the MPS platform lack *in situ* imaging capabilities, limiting the interpretation of certain morphological metrics in real-time. Second, studying hepatocytes alone limits interpretability and certain comparisons with animal models but does allow for a more focused molecular analysis of the drivers of metabolic disease. Co-culture of hepatocytes with islets is an appealing solution to better mimic the normal physiologic insulin delivery ^52^, which exhibits a pulsatility lost in T2D ^53^, but scaling the cell masses and medium volumes to obtain physiological variation is a significant burden. ^54^ Further, only male donors were assessed in this study, which limits the interpretability across sexes, as there are known sexual dimorphisms in metabolic disease. ^55^

In conclusion, our MPS model using primary human hepatocytes exposed to pathophysiological concentration of insulin and nutrients to mimic the hyperinsulinemia, hyperglycemia and increased FFAs to the level observed in T2D subjects recapitulate vital aspects of insulin resistance. This model of insulin resistance offers a means to better understand the molecular mechanisms involved in insulin resistance and provides a baseline for potential therapeutic targets and avenues.

## Acknowledgments

The authors thank Jose L. Cadavid, Rachelle P. Braun, Kairav K. Maniar, and Douglas A. Lauffenburger for their constructive discussions and methodological insights.

## List of Abbreviations

CYP: cytochrome P450
DNL: *de novo* lipogenesis
FFAs: free fatty acids
HGP: hepatocyte glucose production
MPS: microphysiological system
NAFLD: non-alcoholic fatty liver disease
NASH: non-alcoholic steatohepatitis
T2D: type 2 diabetes

