## Supplementary material for "A Microphysiological Model of Progressive Human Hepatic Insulin Resistance": Fig. S

### SUPPLEMENTAL MATERIAL

I SUPPLEMENTAL FIGURES 2

### SUPPLEMENTAL FIGURES

#### Supplemental Figure S1

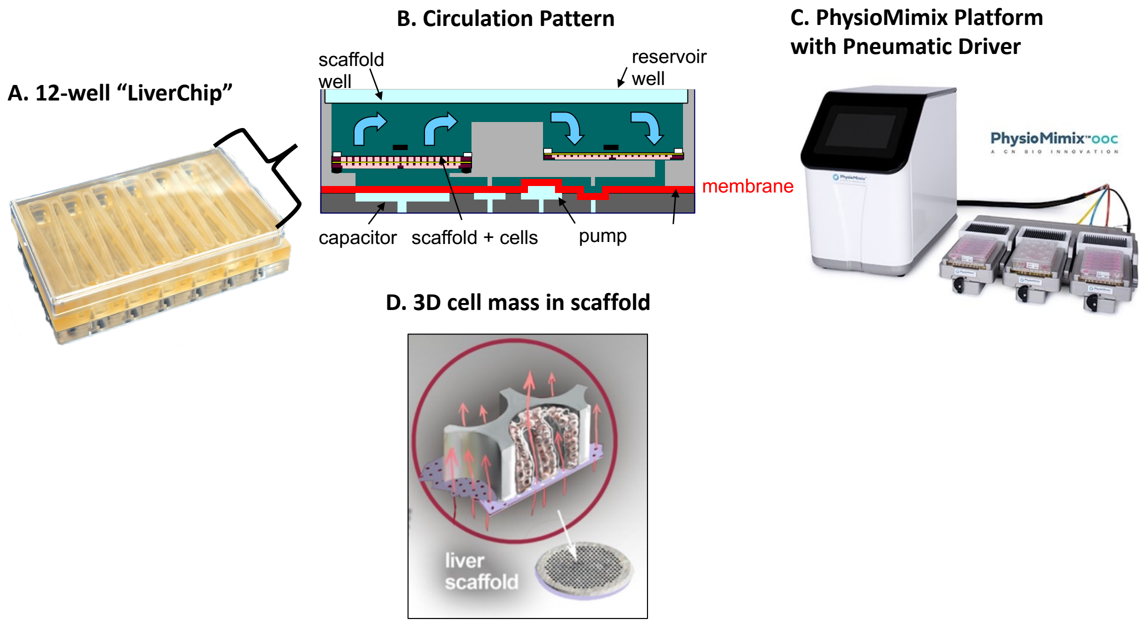


**Supplement Figure S1.** LiverChip design and operation. Each multiwell plate (A) contains 12 identical circulation loops with microfluidic pumps drive (B) driven by pneumatic actuation (C). Cells seeded into the scaffold attach to the walls of the through-holes to form a 3D tissue mass (D).

#### Supplemental Figure S2

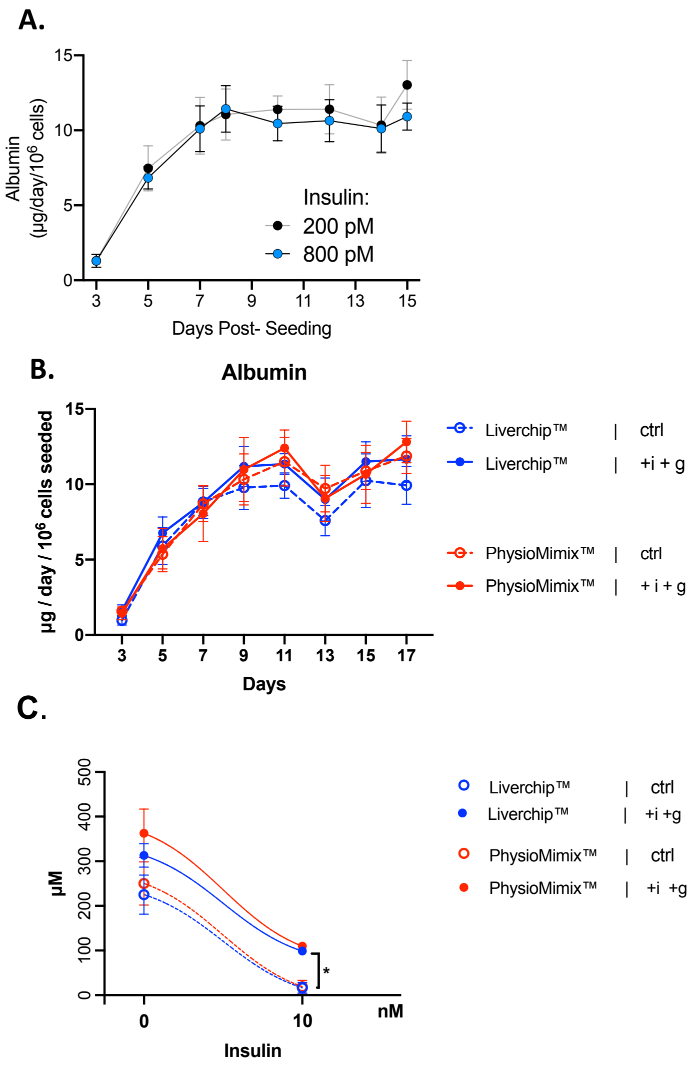


**Figure S2. Albumin production rates stabilize after an initial transient increase and are independent of insulin and nutrient concentration or Bioreactor verson.** **A.** 600,000 hepatocytes (donor SMC) were cultured in the Liverchip™ in either Condition 1 200 pM insulin medium (ctrl) or in Condition 1 + insulin (800 pM) medium. Media change was performed every 48h. The rates of albumin secretion are statistically identical. **B**. 600,000 hepatocytes (donor AQL) were cultured either the Liverchip™ or the PhysioMimix™ in either baseline 200 pM insulin (crtl) medium or in hi-ins (1000 pM) + hi-glucose (i + g) medium. Media change was performed every 48h with the exception of 24h media change between D12 and D13. The rates of albumin secretion are statistically identical in both bioreactor types and both media formulations. (C) 600,000 hepatocytes (donor SMC) were cultured in two bioreactor formats (Liverchip™ vs PhysioMimix™) in two media condtions ( ctrl: physiological | +i +g: hyperinsulinemia + hyperglycemia)(Two-way ANOVA * $p < 0.05$).

**Supplemental Figure S3**

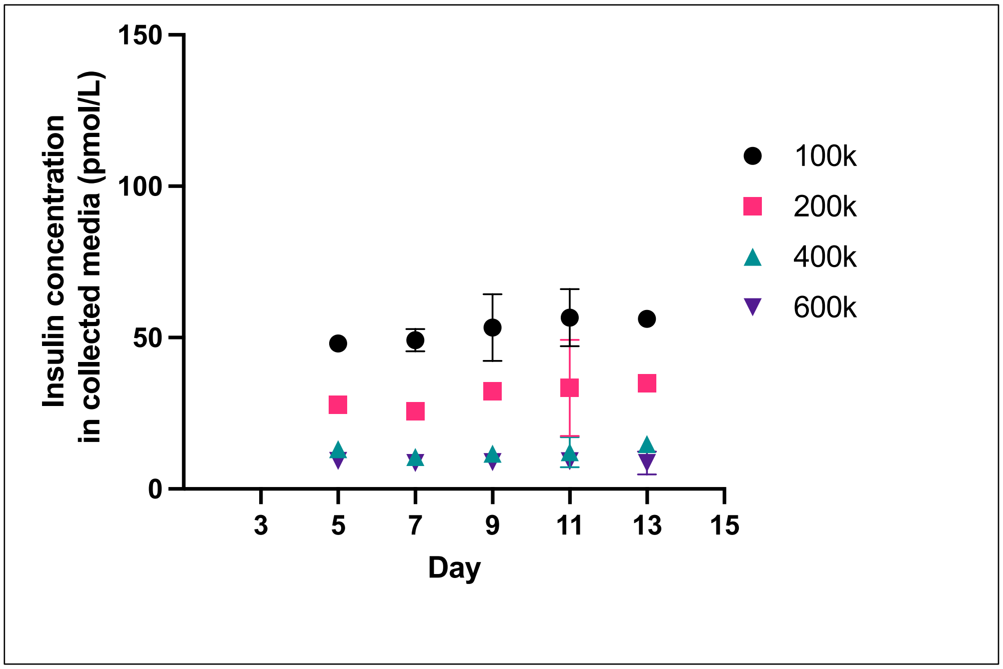


**Figure S3. Insulin depletion from initial 200 pM concentration depends on cell number seeded in PhysioMimix™. A.** Hepatocytes (donor HU2098, ThermoFisher) were cultured in the PhysioMimix™ in baseline 200 pM insulin medium, which was changed every 48 hr and insulin was measured in the collected culture supernate.

**Supplementary Figure S4**

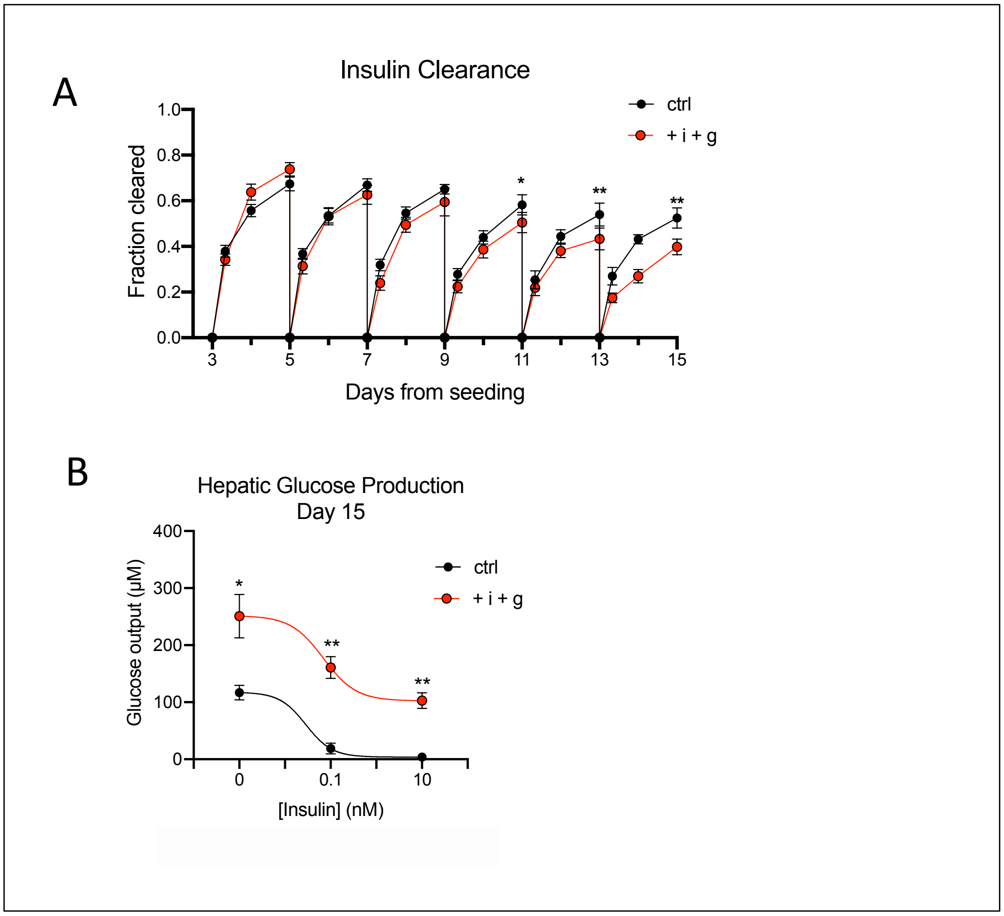


**Figure S4. Hyperinsulinemia and hyperglycemia induce insulin resistance in the 3D perfused liver MPS with donor AQL and alternate bioreactor.** (A) Cells cultured in hi-ins (1000 pM) + hi-glucose (11 mM) conditions (legend: +i +g) gradually lose the ability to clear insulin over 15 days in culture compared to 200 pM insulin baseline medium (legend: ctrl). Two-way ANOVA (* $p < 0.05$, ** $p < 0.01$, n = 6). (B) Cells maintained in hi-ins + hi-glucose show decreased suppression of HGP output upon 24 hour insulin stimulation (interaction term $p < 0.01$, post-hoc Sidak, ** $p < 0.01$, * $p < 0.05$, n = 2). (C) 600,000 hepatocytes (donor SMC) were cultured in two bioreactor formats (Liverchip™ vs PhysioMimix™) in two media condtions ( ctrl: physiological | +i +g: hyperinsulinemia + hyperglycemia)(Two-way ANOVA * $p < 0.05$).

**Supplemental Figure S5.**

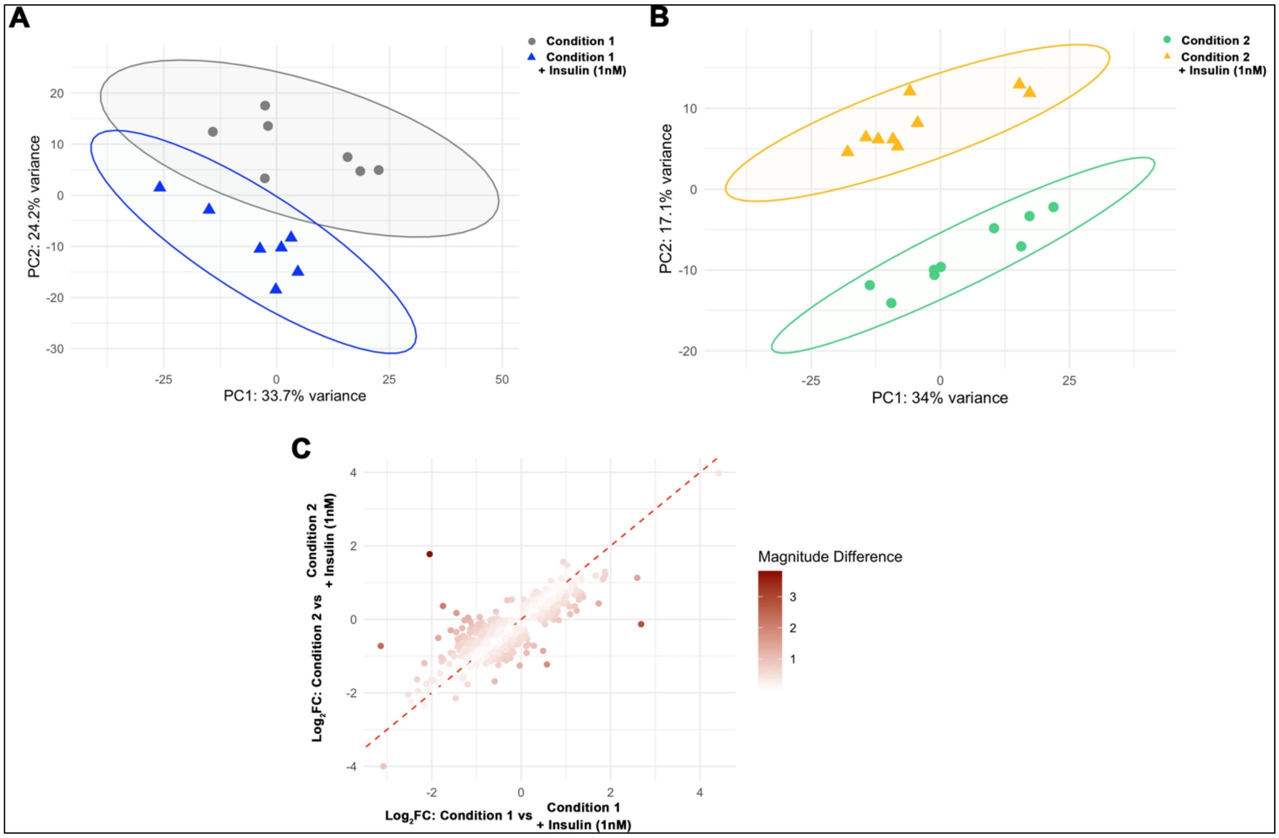


**Figure S5. Hierarchical clustering and magnitude difference of insulin-responsive baseline and disease state samples.** (A,B) Principal component analysis of Condition 1 (left) and Condition 2 (right) samples before and after exposure to 1nM insulin. (C) Magnitude plot of DEGs identified between Condition 1 and Condition 2 samples before and after exposure to 1nM insulin.
